# Effects of a temporary period on pasture on the transcriptomic signature of horses housed in individual boxes

**DOI:** 10.64898/2026.02.09.704829

**Authors:** Aline Foury, Alice Ruet, Nuria Mach, Léa Lansade, Marie-Pierre Moisan

**Author notes:** Corresponding authors who equally contributed to the work Correspondence or.

## Abstract

Pasture access is widely recognized for its welfare, behavioural, and health benefits in horses compared to individual stabling. Our recent studies have shown that even short-term pasture exposure enhances welfare and induces lasting changes in gut microbiota, promoting health-associated bacterial populations. This study investigated the transcriptomic signature of 22 horses following a standardized pasture protocol, with peripheral blood gene expression analysed before pasture access (T0) and three months after returning to individual stabling (T1).

First, using sPLS regression, we correlated gene expression profiles with behavioural welfare indicators measured at T0 and T1. Horses with pasture access exhibited significantly altered blood transcriptome compared to controls, with aggressiveness towards humans emerging as the strongest behavioural correlate of gene expression. Bioinformatic analyses revealed that aggressiveness-associated genes were linked to inflammation, apoptosis, and cell differentiation/growth. Then, horses with pasture access were divided in resilient and non-resilient according to the improvement of their behaviour after time on pasture. Differentially expressed genes post versus pre- pasture, compared between resilient and non-resilient horses for aggressiveness showed that inflammatory signals were downregulated in both subgroups, with a more pronounced effect in resilient horses. This suggests that resilient individuals are better equipped to modulate inflammatory responses in low-stress environments like pasture. Surprisingly, for unresponsiveness to the environment—a trait linked to depressive-like states—resilient horses displayed increased inflammatory signaling (e.g., IRF and CD40 activation) post-pasture, while non-resilient horses showed activation of anti-inflammatory PPAR signaling. Notably, non-resilient horses exhibited molecular signatures associated with organismal death, morbidity, and growth failure, indicating maladaptive physiological states. In contrast, resilient horses demonstrated activation of growth-related pathways (e.g., BMP2 and BMPR1A), suggesting a shift toward anabolic and developmental processes. This study underscores the behavioural and molecular benefits of pasture access for horses, particularly in reducing aggressiveness and inflammatory signaling in resilient individuals. The findings highlight the complex interplay between behaviour, inflammation, and resilience, with pasture access promoting adaptive physiological and molecular responses. Further research is needed to elucidate the long-term implications of these transcriptomic changes and their broader relevance to equine welfare.

## Introduction

Over the past two decades, improving horse welfare through better housing conditions has emerged as a major concern (de Oliveira and Aurich, 2021; Gehlen et al., 2021; Popescu et al., 2019; Ruet et al., 2019). Numerous studies have consistently demonstrated that individual housing negatively impacts equine health and behaviour. These adverse effects are mostly induced by restricted opportunities for social interactions (Søndergaard et al., 2011), limited freedom of movement (Houpt et al., 2001), and the absence of continuous grazing (Harris, 1999)–factors identified as fundamental behavioural needs in a recent meta-analysis (Krueger et al., 2021). Horses kept permanently on pasture with conspecifics typically display a more natural behavioural repertoire, particularly in feeding patterns and social interactions (Christensen et al., 2002; King et al., 2013), and they tend to show fewer health problems (Yngvesson et al., 2019). This evidence suggests that groups living on pasture support a higher welfare state compared with individual stabling (Hartmann et al., 2012). However, it remains unclear whether short-term access to pasture confers similar benefits for horses that are normally housed alone.

To address this question, we monitored a population of horses accustomed to year-round individual stabling before and after a 1.5-month period on pasture, and compared them with a control group of individually housed horses that remained in their boxes. At the behavioural level, welfare of horses on pasture improved but the beneficial effects did not last when horses return to individual boxes (Ruet et al., 2020). Fecal microbiota analyses showed that pasture exposure induced an increase in butyrate-producing bacteria that lasted 1 month after the return to individual boxes, which may have promoted beneficial effects on health and welfare (Mach et al., 2021).

To complete these data, we conducted blood transcriptome analyses, and we performed differential and regression analyses of gene expression to identify molecular pathways associated with behavioural indicators of welfare impairment in sport horses, as defined by Ruet et al. (2022). Our results showed that access to pasture didn’t have the same effect on the different behavioural indicators and that aggressiveness toward humans is the behavioural trait most strongly correlated with alterations in blood gene expression. The associated molecular pathways are predominantly linked to systemic inflammatory processes. Time in pasture has opposite effects depending on the behavioural indicator. When horses that went on pasture were divided in resilient and non-resilient according to the improvement of their behaviour, inflammatory processes were more activated for non-resilient horses for aggressiveness toward humans whereas less activated for non-resilient horses for unresponsiveness to the environment.

## Material and methods

### Animals

The study included 22 sport horses that had lived in individual boxes since they were three years of age. Prior to this, these horses had lived outside on pasture in groups on their breeding farms. Horses were recruited from a cohort of 60 individuals housed at the French National Riding School, in Saumur, France (Ruet et al., 2020).

The selected horses were randomly split into two groups: 8 horses (6 geldings and 2 mares) aged 9.88 ± 3.83 years (mean ± SD) were kept in individual boxes (Control group) whereas 14 horses (8 geldings and 6 mares) aged 9.57 ± 2.82 years were moved to pasture (Pasture group). Horses in the Control group were permanently housed in individual boxes of 9 m^2^. All horses had visual contact with conspecifics and reduced tactile contacts through a grilled window on the wall between two boxes. They were trained for sport purposes six days out of seven. Horses of the Pasture group spent 41.7 ± 16.8 days on pasture between August and September and then returned to their boxes. The pastures were located 5 ± 1.5 km from the riding school. The average surface area of the pastures was 5.02 ± 0.4 ha, which was much larger than the minimum recommended surface area ensuring a low level of aggression among horses (0.03 ha per horse; Flauger and Krueger, 2013).

### Behavioural assessment

Four independent behavioural indicators reflecting compromised welfare state were recorded during the experiment: aggressive behaviours towards humans, unresponsiveness to the environment, alert posture and stereotypies (see Ruet et al., 2022 for the description of these indicators). The two groups were studied during two different periods: just before pasture (T0) and 3 months after returning to the box after the pasturing period (T1). For each period, the horses were observed over five consecutive days using the scan sampling method (Altmann, 1974). Per period, behavioural observations were carried out during 10 sessions of 90 min each. Thirteen scans per horse were recorded per session. Details of the methodology applied are described in Ruet et al. (2020).

### Blood RNA extraction

At T0 and T1, venous blood (10 ml) was collected once between 09:00 and 12:00 from the horses using EDTA tubes (BD Vacutainer®). In order to stabilize intracellular RNA, 0.8 ml of venous blood was transferred to a tube containing 0.8 ml of Lysis Buffer DL (Macherey-Nagel, Düren, Germany). After homogenization, the samples were maintained at - 20 °C until RNA extraction. Total RNA was extracted using the NucleoSpin® RNA Blood kit (Macherey-Nagel, Düren, Germany). The manufacturer’s protocol was modified to obtain sufficient ARN quantities for the microarray gene expression analyses: extraction was carried out from 1.6 ml of the mixed blood and Lysis Buffer DL. Twenty μl of proteinase K was then added to complete blood lysis. To adjust RNA binding conditions, 0.8 ml of 70% ethanol was added. The following steps were not modified, except the elution that was carried out with 40 μl of RNase-free water. Total RNA quality was assessed using RNA Pico chips on a Bioanalyzer 2100 (Agilent, Boeblingen, Germany), and its concentration was measured on a NanoDrop One Spectrophotometer (ThermoScientific, Illkirch, France).

### Microarray gene expression analyses

Gene expression profiles were performed at the GeT-TRiX facility (GénoToul, Génopole Toulouse Midi-Pyrénées) using Agilent Sureprint G3 Horse_60 K_2016_01_22 021322 GE microarrays (8 × 60 K, design AMADID 081421) following the manufacturer’s instructions. For each sample, Cyanine-3 (Cy3) labelled cRNA was prepared from 50 ng of total RNA using the One-Color Quick Amp Labeling kit (Agilent Technologies) according to the manufacturer’s instructions, followed by Agencourt RNAClean XP (Agencourt Bioscience Corporation, Beverly, Massachusetts). Dye incorporation and cRNA yield were checked using Dropsense™ 96 UV/VIS droplet reader (Trinean, Belgium). A total of 600 ng of Cy3-labelled cRNA were hybridized on the microarray slides following the manufacturer’s instructions. Immediately after washing, the slides were scanned on Agilent G2505C Microarray Scanner using Agilent Scan Control A.8.5.1 software. The fluorescence signal was extracted using Agilent Feature Extraction software v10.10.1.1 with default parameters. Microarray data and experimental details are available in NCBI’s Gene Expression Omnibus (Barrett and Edgar, 2006) and are accessible through GEO Series accession number GSE310717. https://www.ncbi.nlm.nih.gov/geo/query/acc.cgi?acc=GSE310717

### Impact of blood cell proportions in gene expression profiles

We assessed whether the observed gene expression changes in the different analyses (Control vs. Pasture at T0 and T1) were related to changes in cell proportions in the blood samples using the Celltype Computational Differential Estimation CellCODE R package (Chikina et al., 2015).

### Statistical analyses

The percentage of scans for each behavioural indicator per period was calculated from the total number of observations per horse. Behavioural expression showed a great variation before and after pasture, and as we wanted to characterize the differences in gene expression between horses that retained the benefit of pasturing from those that did not, we split pasture horses into 2 groups: Resilient horses (R) were horses that reduced their behavioural expression more than 1% after pasture, the other pasture horses were considered Non-Resilient (NR). Paired non-parametric t-test followed by Wilcoxon matched-pairs signed rank tests were used to compare the percentages of scans for the behavioural indicators before and after pasture. Statistical analyses were performed using GraphPad Prism (version 10.5.0) with a significance level of P ≤ 0.05.

Microarray data were analysed using the R software (R version 4.1.3; R Core Team, 2021) and Bioconductor packages (http://www.bioconductor.org, v.3.15; Huber et al., 2015) as described in GEO accession GSE310717. Raw data (median signal intensity) were filtered, log2 transformed, corrected for batch effects (microarray washing bath and labelling serials), and quantile normalized (Bolstad et al., 2003).

For the annotation of the Agilent chip probes, we used the annotation provided by Agilent and for the non-annotated probes, the annotation was performed using Nucleotide BLAST (https://blast.ncbi.nlm.nih.gov/). BLASTs were performed with the following parameters: Organism = Equus caballus; minimum cover = 95%, minimum identity = 95%.

The relationships between gene expression and welfare indicators were analysed using the R package mixOmics v.6.18.1 (Rohart et al., 2017). First, PCA and Sparse Partial Least Square (sPLS) analysis were performed to explore the data set structure and characterize the correlation between gene and behavioural indicators expressions. The functions of the genes selected through sPLS were annotated using GeneCards (http://www.genecards.org/), and each list of genes was analysed using Ingenuity Pathway Analysis (IPA, QIAGEN Inc., https://digitalinsights.qiagen.com/).

For the differential analyses, a model was fitted using the limma lmFit function (v.3.50.3, Ritchie et al., 2015) considering animals as a blocking factor to account for repeated measures, using the duplicateCorrelation function. Pair-wise comparisons between biological conditions were applied using specific contrasts. A correction for multiple testing was applied using the Benjamini-Hochberg procedure (BH, Benjamini and Hochberg, 1995) to control the False Discovery Rate (FDR). Probes with an FDR ≤ 0.05 and showing a difference ≥ 20% in mean expression levels between biological conditions were considered to be differentially expressed between conditions. As for sPLS, gene lists were analysed using Ingenuity Pathway Analysis (IPA, QIAGEN Inc., https://digitalinsights.qiagen.com/).

## Results

### Evolution of the behavioural indicators during pasture

Four independent behavioural indicators reflecting compromised welfare state were recorded pre-(T0) and post-pasture (T1): aggressive behaviours towards humans, unresponsiveness to the environment, alert postures, and stereotypies (Table 1). Figure 1 shows the evolution of the behavioural indicators between T0 and T1 for each horse. Variability in behavioural expression between T0 and T1 is important, particularly for Pasture horses. More generally, apart from unresponsiveness to the environment, behaviours were rarely expressed for either group, at T0 or T1: the majority of values are therefore equal to 0. This is why none of the analyses performed between T0 and T1 yielded significant results.

**Table 1.** Percentage of scans recorded for aggressive behaviours towards humans, unresponsiveness to the environment, alert postures and stereotypies before pasture (T0) and 3 months after returning to the box after the pasturing period (T1) (mean ± SD). Scans were also recorded for the Control group (box only) at the same periods. Before and after pasture comparisons: Wilcoxon matched-pairs signed rank tests. ns=non-significant.

|  | n | Control<br>8 | Pasture<br>14 |
| --- | --- | --- | --- |
| Aggressiveness | T0 | 0.50 ± 1.40 | 4.75 ± 7.09 |
| | T1 | 0.25 ± 0.71 | 2.91 ± 3.99 |
|  | P | ns | ns |
| Unresponsiveness | T0 | 2.78 ± 2.74 | 3.42 ± 3.56 |
| | T1 | 2.99 ± 2.56 | 2.58 ± 4.97 |
|  | P | ns | ns |
| Alert | T0 | 0.36 ± 0.51 | 1.02 ± 1.80 |
| | T1 | 0.15 ± 0.30 | 1.60 ± 2.84 |
|  | P | ns | ns |
| Stereotypies | T0 | 0.22 ± 0.62 | 0.31 ± 0.93 |
| | T1 | 0.28 ± 0.61 | 0.93 ± 1.80 |
|  | P | ns | ns |

**Figure 1.**
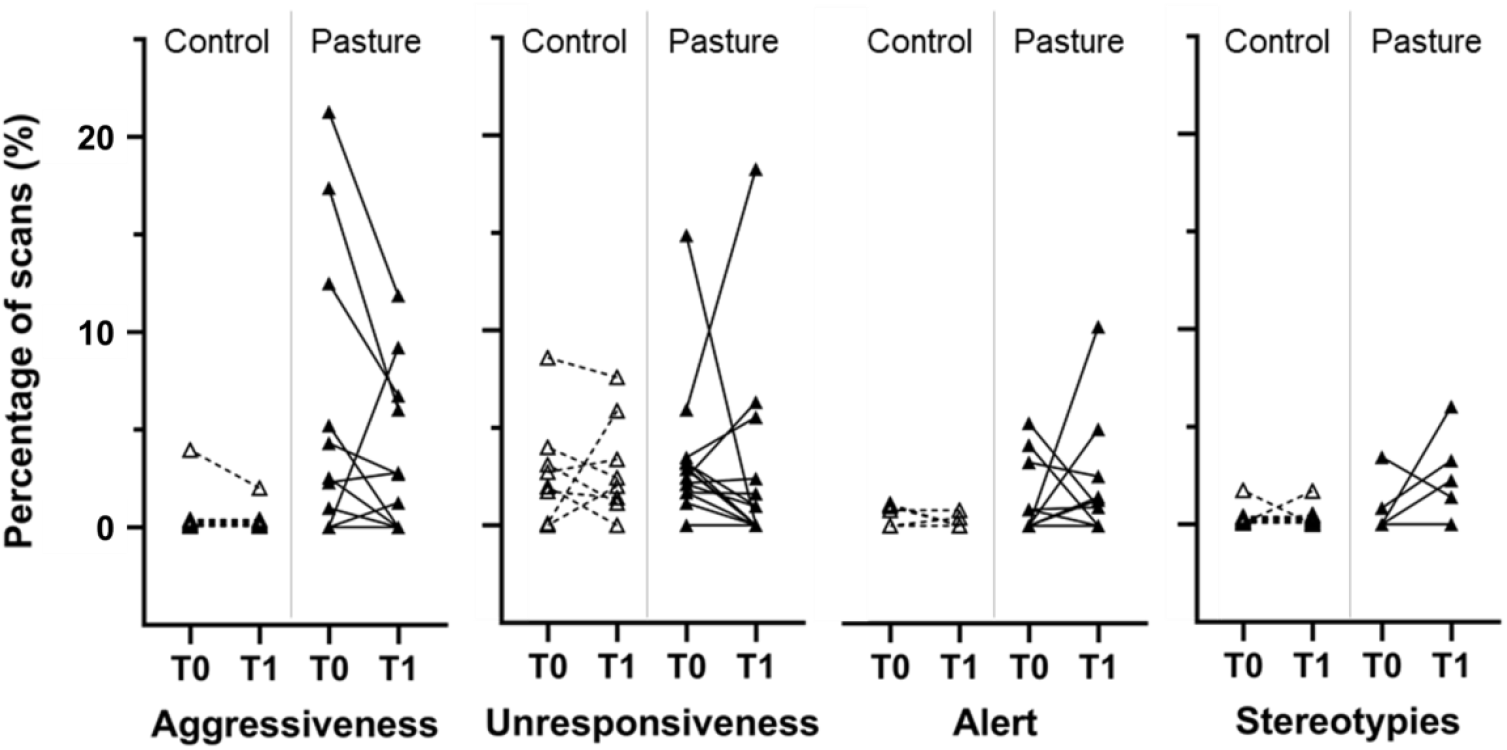
Percentage of scans recorded for aggressive behaviours towards humans, unresponsiveness to the environment, alert postures and stereotypies before pasture (T0) and 3 months after returning to the box after the pasturing period (T1). Scans were also recorded for the Control group (box only) at the same periods.

### Gene expression analyses

We found no significant changes in the proportion of immune cell subtypes in the different analyses using CellCODE (Table S1).

#### Exploratory regression analysis between peripheral blood transcriptome and behavioural indicators

Preliminary PCA of the transcriptomic dataset (Figure S1) showed that gene expression is largely influenced by the sampling period, especially in Pasture horses. A sparse Partial Least Square (sPLS) analysis was then performed to explore the data set structure and characterize the correlation between gene and behavioural indicators expressions, and identify the genes that contributed the most to explaining differences between contrasted horses for each behavioural indicator. Very few horses expressed stereotypies at T0 (1 Control horse and 2 Pasture horses) or at T1 (3 Control horses and 4 Pasture horses). As sPLS could not be performed on a dataset with a majority of data equal to 0, this indicator was excluded from the analysis. The analysis included the 2000 genes that were the most relevant for the model and the three behavioural indicators left. The projection of the samples into the space spanned by the averaged components of gene and behavioural indicators expression datasets shows, as expected, that time has a much stronger effect on gene expression for Pasture horses than for Control Horses (Figure 2).

**Figure 2.**
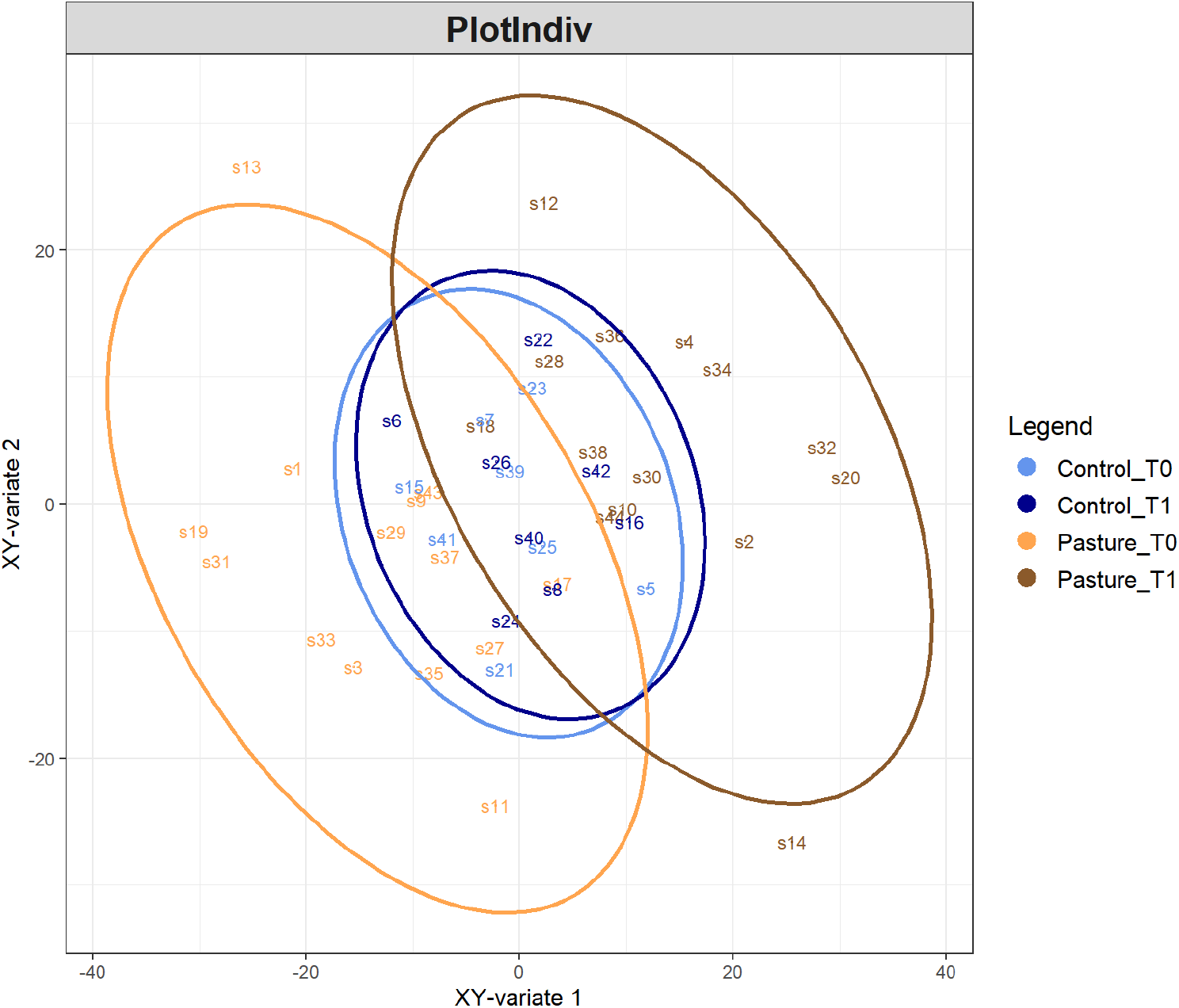
sPLS sample plot. Samples are projected into the space spanned by the averaged components of both gene expression and behavioural indicators datasets.

Correlation levels between gene expression and behavioural indicator variables are presented in Figure 3. Genes ‘highest levels of correlation were found with aggressiveness towards humans. Arising from these results, we focused our downstream analysis on the “aggressiveness towards humans” trait. We identified 825 genes correlated with aggressiveness (absolute correlation value ≥ 0.5): 414 were positively and 411 negatively correlated. We then used Ingenuity Pathway Analysis (IPA) software to better understand positively correlated genes’ functional implications and reveal the molecular pathways and upregulators underpinning this gene set. Figure S2 shows that the over-represented biological process terms were strongly associated with inflammatory response, e.g., interleukins IL-1A, IL-1N, IL6, IL18, and genes NFKB1 (Nuclear Factor k B1), TLR4 (Toll-like Receptor 4), TNF (Tumour Necrosis Factor) expression, leukopoiesis, lymphopoiesis. Using the IPA software’s upstream regulator analysis, we identified 70 upstream regulators with p-values < 0.0001 (Table S2). These regulators were related to the onset of inflammation, apoptosis, and cell differentiation/growth. The list of estimated up-regulators is summarised in Figure 4 through a CIRCOS plot.

**Figure 3.**
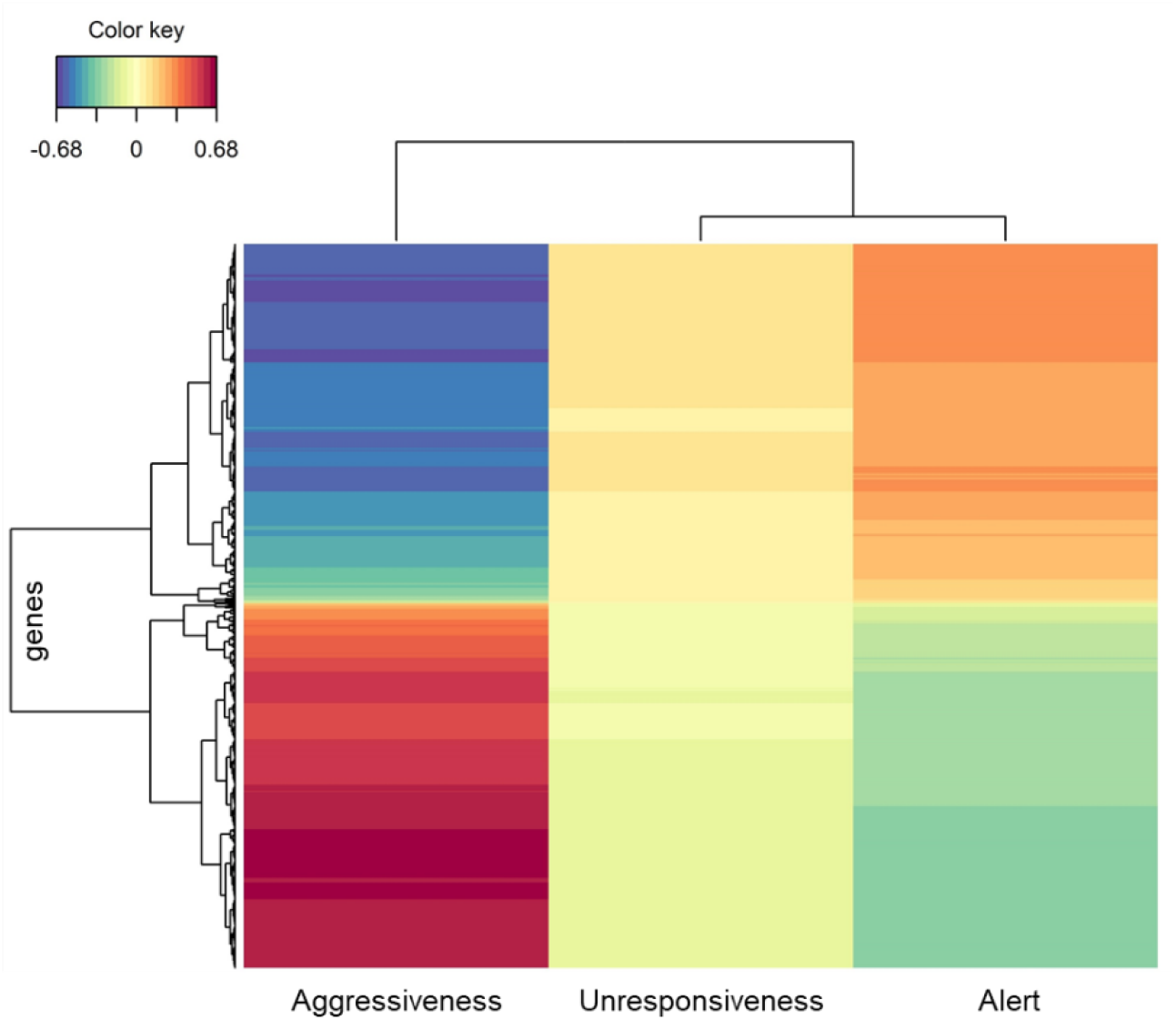
Heatmap from sPLS analysis. The plot displays the correlation levels between gene expression and the behavioural indicator variables. Genes are clustered according to the direction and the intensity of the correlation with behavioural indicators. The most positive and negative correlations are represented in dark red and dark blue, respectively, as indicated in the colour key legend.

**Figure 4.**
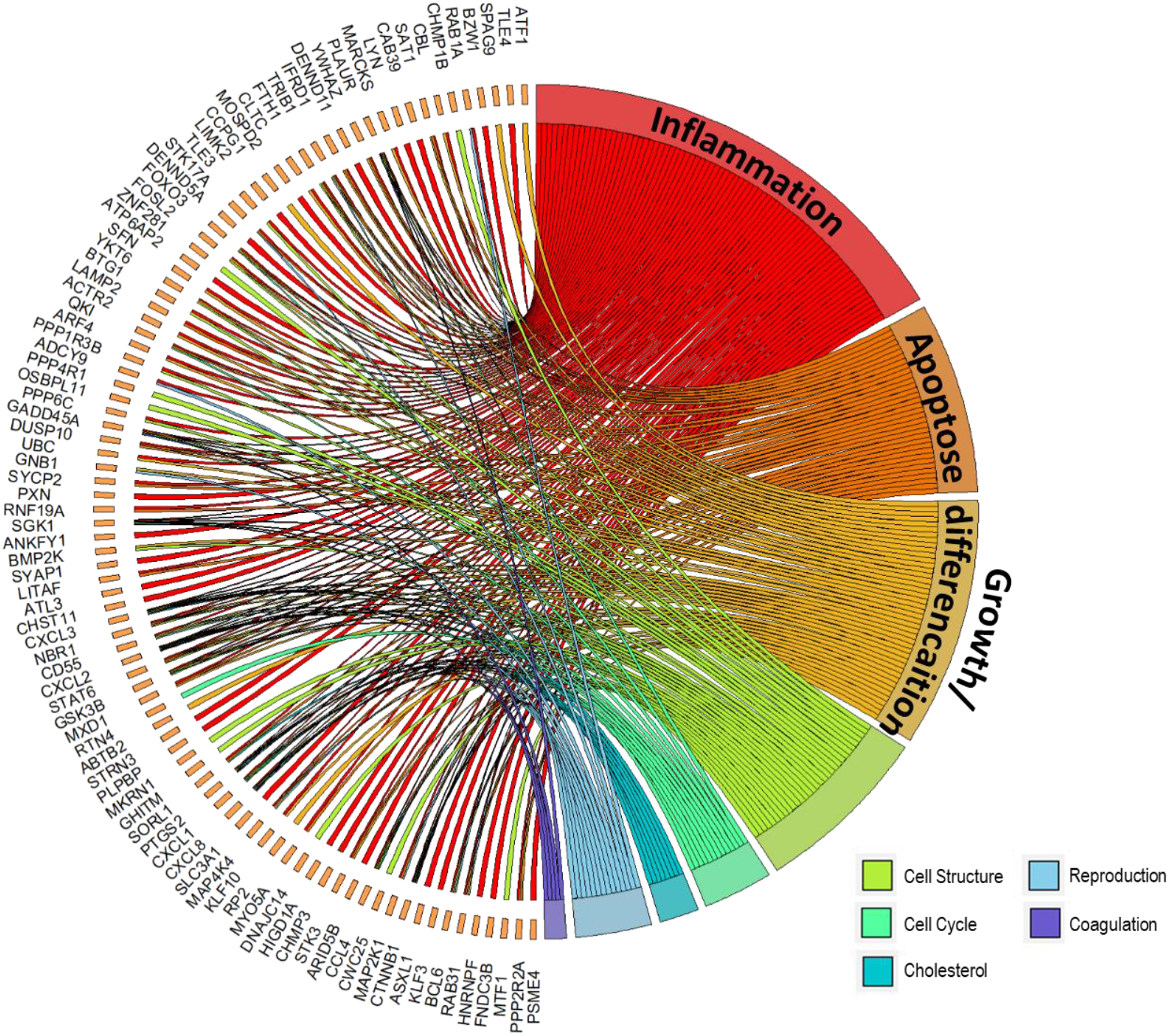
CIRCOS plot of main upregulators and their target genes positively correlated to aggressiveness towards humans.

#### Differential analyses between resilient and non-resilient horses

To characterize the differences in gene expression between horses that retained the benefit of pasturing and those that did not, we divided the Pasture horses into two groups: Resilient horses (R) that displayed a reduction in their behavioural expression of more than 1% after pasture, and Non-Resilient horses (NR) that did not show this decrease in behavioural expression. The > 1% decrease in behavioural expression is used as a quantitative proxy for the horse’s ability to recover, regulate, and adapt – key components of resilience. Thus, resilient individuals exhibit flexible, context-appropriate modulation of behaviour following a challenge or environmental change, whereas non-resilient individuals show persistently heightened behavioural responses. For alert postures, the number of R and NR horses was so unbalanced that we were unable to perform statistical analyses (2 R horses vs. 12 NR horses). Consequently, statistical analysis between R and NR horses was only carried out for aggressive behaviours towards humans and unresponsiveness to the environment. The percentage of scans recorded for those two behavioural indicators at T0 and T1 is presented in Table 2. Because of our classification, there is a significant difference in the frequency of behaviour expression between T0 and T1 for R horses only.

**Table 2.**
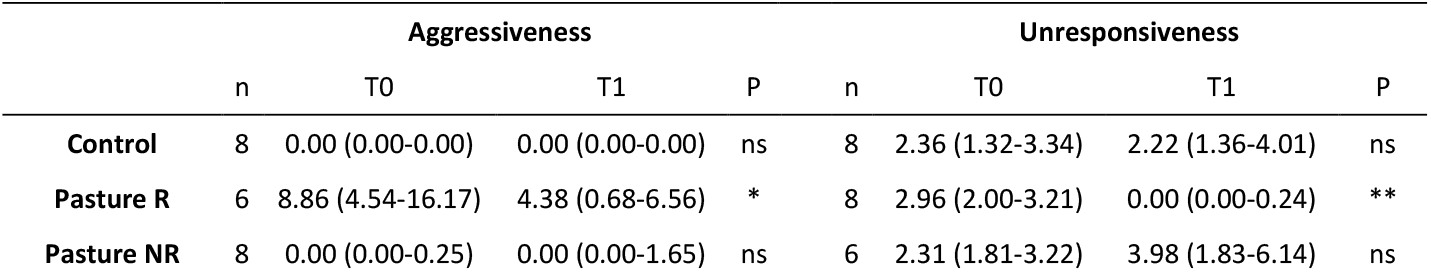
Percentage of scans recorded for aggressive behaviours towards humans and unresponsiveness to the environment before pasture (T0) and 3 months after returning to the box after the pasturing period (T1) (median and interquartile range). Scans were also recorded for the Control group (box only) at the same periods. Pasture horses were divided into two groups: Resilient horses (R) were horses that reduced their behavioural expression after pasture more than 1%, and Non-Resilient horses (NR) were horses that did not. Before and after pasture comparisons: Wilcoxon matched-pairs signed rank tests. ns=non-significant; *P ≤ 0.05; **P ≤ 0.01.

|  | Aggressiveness |  |  |  | Unresponsiveness |  |  |  |
| --- | --- | --- | --- | --- | --- | --- | --- | --- |
|  | n | T0 | T1 | P | n | T0 | T1 | P |
| **Control** | 8 | 0.00 (0.00-0.00) | 0.00 (0.00-0.00) | ns | 8 | 2.36 (1.32-3.34) | 2.22 (1.36-4.01) | ns |
| **Pasture R** | 6 | 8.86 (4.54-16.17) | 4.38 (0.68-6.56) | * | 8 | 2.96 (2.00-3.21) | 0.00 (0.00-0.24) | ** |
| **Pasture NR** | 8 | 0.00 (0.00-0.25) | 0.00 (0.00-1.65) | ns | 6 | 2.31 (1.81-3.22) | 3.98 (1.83-6.14) | ns |

#### Resilience effect on aggressiveness towards humans

Differential analyses were performed to investigate the effect of pasture on gene expression in both resilient (R.T1-R.T0) and non-resilient (NR.T1-NR.T0) horses for aggressiveness towards humans. At an FDR ≤ 0.05 and a fold change ≥ 20%, we identified 1903 differentially expressed genes in resilient horses and 1283 in non-resilient horses. The IPA software’s comparative analysis was used to compare the effect of pasture in R and NR horses, revealing differences in the canonical pathways, upstream regulators, and biological functions in which those differentially expressed genes are involved (Figure 5).

**Figure 5.**
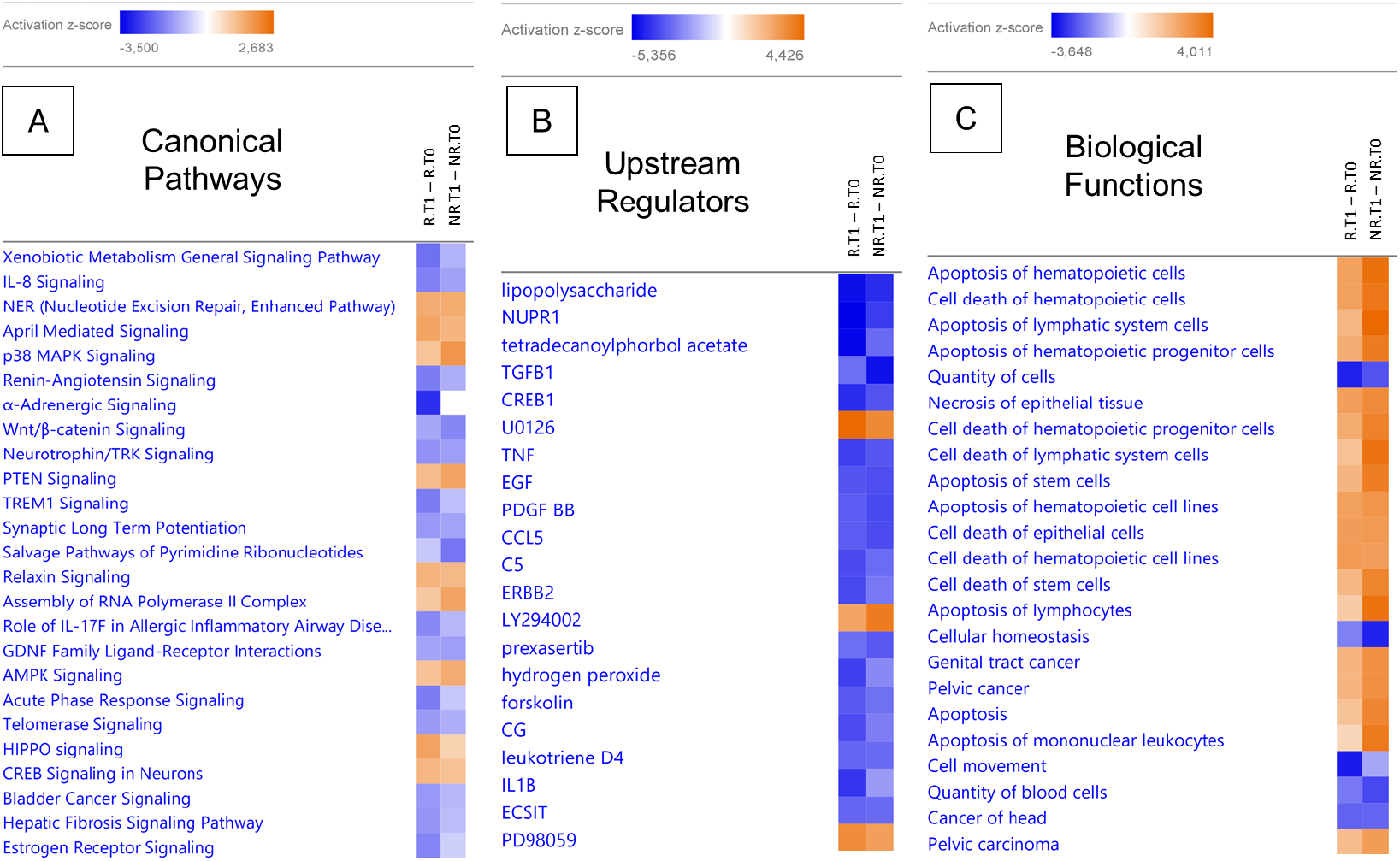
Comparative analysis between time in pasture effect differential analyses for Resilient (R) and non-resilient (NR) horses for aggressiveness towards humans, generated through the use of QIAGEN IPA (QIAGEN Inc., https://digitalinsights.qiagen.com/IPA): Canonical Pathways (A), Upstream regulators (B), and Biological functions (C). As indicated in the colour key legend, activation z-scores were the most negative for dark blue and the most positive for dark orange.

The comparative analyses show that canonical pathways (Figure 5A) related to inflammation were more down-regulated for R horses: IL-8 (Interleukin 8) signaling, αβ-adrenergic signaling, TREM1 (Triggering Receptor Expressed on Myeloid cells) signaling, Role of IL-7F in allergic inflammatory airway disease, and Acute Phase Response signaling. Similarly, inflammatory upstream regulators (Figure 5B) (CREB (cAMP Response Element-Binding protein), TNF (Tumor Necrosis Factor), and IL1β) and many biological functions related to apoptosis are less active in R horses (Figure 5C).

When we analysed R and NR horses using differential analysis, we found no differentially expressed genes at T1 with FDR≤0.05. To better understand this result, we examined the level of aggressiveness of each R and NR horse at T0 and T1 (Figure S3). The R horses for aggressiveness had the highest levels of aggressiveness of the whole population at T0, which was drastically reduced at T1, while the NR horses displayed low or moderate aggressiveness at both T0 and T1. These data explain why no differentially expressed genes are detected between R and NR at T1.

#### Resilience effect on unresponsiveness to the environment

Differential analyses with an FDR ≤ 0.05 and a fold change ≥ 20% showed significant results between T0 and T1, with 1357 differentially expressed genes for R horses and 2043 for NR horses. As for aggressiveness, a comparative analysis using IPA software was performed (Figure 6).

**Figure 6.**
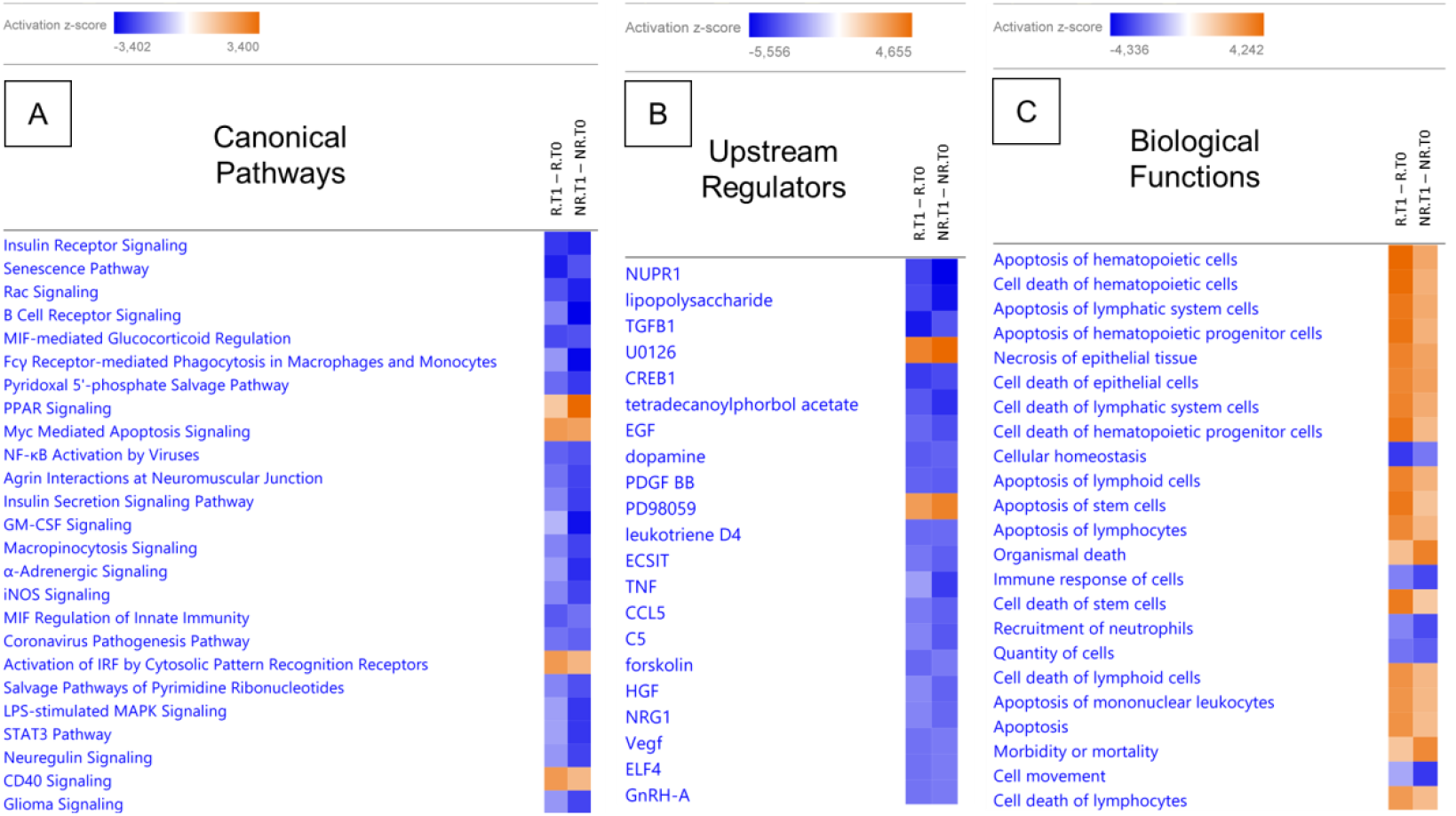
Comparative analysis between time in pasture effect differential analyses for Resilient (R) and non-resilient (NR) horses for unresponsiveness to the environment, generated through the use of QIAGEN IPA (QIAGEN Inc., https://digitalinsights.qiagen.com/IPA): Canonical Pathways (A), Upstream regulators (B) and Biological functions (C). As indicated in the colour key legend, activation z-scores were the most negative for dark blue and the most positive for dark orange.

The comparative analyses show that B cell receptor signaling, Fcγ Receptor-mediated phagocytosis, GM-CSF (Granulocyte-macrophage colony-stimulating factor) signaling, α-Adrenergic signaling, and LPS(LipoPolySaccharide)-stimulated MAPK (Mitogen-Activated Protein Kinase signaling pathways were more down-regulated in NR than R horses, and PPAR (Peroxisome Proliferator–Activated Receptors) signaling was more activated, while CD40(Cluster of Differentiation 40) and IRF (Interferon Regulatory Factor) signaling were more activated in R than NR horses (Figure 6A). Inflammatory upstream regulator CREB1 was more down-regulated in R horses, whereas TNF was more down-regulated in NR horses (Figure 6B). Many biological functions related to apoptosis are less active in NR horses, but organismal death, morbidity, and mortality were more active in the same NR horses (Figure 6C).

Differential analysis also revealed an effect of resilience at T1 with 449 differentially expressed genes between R and NR horses. The main biological process in which those differentially expressed genes are involved is growth which is activated for R horses and inhibited for NR horses (Figure 7).

**Figure 7.**
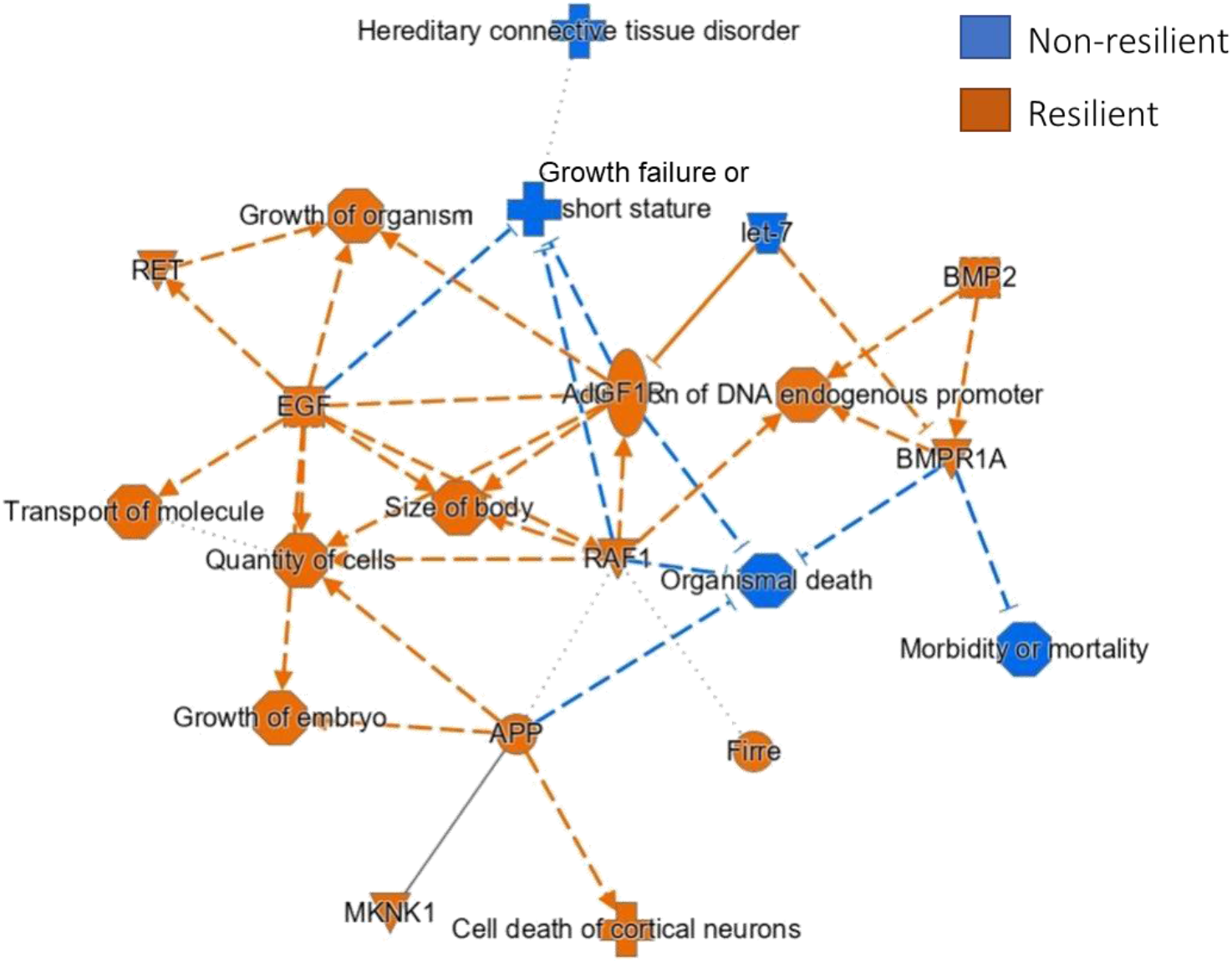
Graphical Summary of differentially expressed genes (FDR ≤ 0.05; difference ≥ 20% in mean expression levels between Resilient and Non-Resilient horses) at T1, for unresponsiveness to the environment, generated through the use of QIAGEN IPA (QIAGEN Inc., https://digitalinsights.qiagen.com/IPA).

## Discussion

Keeping horses on pasture, when managed appropriately, provides clear welfare, behavioural, and health advantages over stabling. Even short-term pasture access can improve welfare, though adaptation periods are needed, and behavioural benefits diminish when horses return to stabling (Ruet et al., 2020). The gut microbiota of horses kept in pasture for 1.5 months shows an increase in health-promoting bacteria that persists 1 month after the horses return to their individual boxes (Mach et al., 2021). In this study, we analysed the transcriptomic signature of 22 horses that followed the same protocol, with peripheral blood gene expression measured before pasture access (T0) and 3 months after the horses returned to their individual box (T1).

As a first exploration of the peripheral blood gene expression profiles of the 22 horses, we performed an sPLS regression analysis with the variables related to behavioural indicators of welfare, just before pasture and 3 months after their return from pasture. Not surprisingly, horses that spent time in the pasture had their blood transcriptome much more affected than the control horses that stayed in their box. The strongest indicator correlated with gene expression was aggressiveness towards humans. Bioinformatic analyses of the aggressiveness-correlated genes revealed that the upstream regulators were related to the onset of inflammation, apoptosis, and cell differentiation/growth. This pattern suggests that aggressiveness in horses may be underpinned by a molecular landscape reflecting heightened immune vigilance and tissue-remodelling dynamics, possibly linked to the increased risk of physical confrontation or injury inherent to aggressive interactions. We found the same results in a previous study on 45 sport horses kept in individual boxes (Foury et al., 2023). The highest correlation of genes expression with aggressiveness is thought to be due to the less labile aspect of aggressiveness behaviour compared to unresponsiveness to the environment or alert postures (Foury et al., 2023). While vigilance or environmental reactivity may fluctuate rapidly in response to contextual stimuli, aggressiveness likely represent a more consistent individual characteristic supported by long-term physiological and molecular profiles. Increased inflammation related to aggressiveness was found by others in mice (Malki et al., 2014) and zebrafish (Reichmann et al., 2022) transcriptomic analyses. In horses subjected to individual stabling, stress-related changes in immune cell profiles and increased stress hormone (cortisol) levels were found, which are known to modulate inflammation. These changes were accompanied by disturbed behaviours, including increased aggression, highlighting the role of environmental stressors in driving both inflammation and aggressive responses (Schmucker et al., 2022). The hypothesis posits that upregulating immune functions during and after aggressive encounters—where physical injury is likely—is advantageous for the organism.

When we compared the effect of time in pasture on gene expression in resilient and non-resilient horses within the Pasture group, we found that inflammatory signals are down-regulated in both subgroups, but more so in the resilient horses, suggesting that these individuals are more capable of modulating and dampening inflammatory pathways in response to a positive, low-stress environment such as pasture access. These data confirmed that inflammation is associated with the level of aggressiveness in horses. Three months after their return to the box, the resilient horses were no longer different from the non-resilient ones regarding transcriptomic signature. This can be explained by the fact that the resilient horses displayed more aggressiveness before going to pasture (T0) compared to the non-resilient ones.

Regarding unresponsiveness to the environment, signaling related to inflammation seems more activated in the resilient subgroup of horses that spent time on pasture. Indeed, IRF as well as CD40 signaling were activated after time in pasture in resilient horses, while PPAR signaling that has anti-inflammatory effects, is up-regulated in non-resilient horses after pasture. Unresponsiveness to the environment is thought to reflect a depressive-like state in horses (Fureix et al., 2012). Depressive symptoms are known to be associated with increased inflammation, so the data observed here do not fit with resilient horses showing higher inflammation. Among the resilient horses for unresponsiveness, two of them display high aggressiveness, which is strongly associated with inflammation as discussed above. Further analyses are required to conclude whether this observation may explain the increased inflammatory signals in the resilient group of horses for unresponsiveness.

Besides decreased inflammatory signals, the comparative analysis shows that the non-resilient horses for unresponsiveness display an activation of organismal death, morbidity, and mortality, suggesting a maladaptive or costly physiological state, in which unresponsiveness is accompanied by molecular signatures reflecting chronic stress, inefficient coping, or a failure to appropriately resolve inflammatory and metabolic challenges. This result is confirmed in the differential analysis comparing resilient and non-resilient horses after pasture (T1). Indeed, organismal death and mortality occur in NR horses alongside growth failure or short stature. Regarding the R horses, they display an activation of functions such as growth of embryo, growth of organism, size of the body, as well as an activation of BMP2 (Bone Morphogenetic Protein 2) growth factor and its type 1A receptor BMPR1A that play an important role in the development of bone and cartilage. In growing animals, such engagement of developmental or remodelling program associated with decreased unresponsiveness to the environment looks beneficial for health. Taken together, these findings support the hypothesis that behavioural resilience is mirrored at the molecular level by a shift towards anabolic, developmental and remodelling pathways, whereas non-resilience is associated with catabolic, degenerative and risk-laden signatures.

In conclusion, this study shows that spending time on pasture is beneficial for some horses, both at the behavioural level and in the blood transcriptome. Aggressiveness towards humans is the behavioural trait most strongly correlated with gene expression and linked to inflammation, apoptosis, and growth/differentiation. Short-term time in pasture was beneficial for some of the horses (R horses), which displayed reduced aggressiveness and decreased inflammatory signaling. Horses that improved their unresponsiveness to the environment display a transcriptomic signature associated with elevated inflammation after time in pasture, but also with increased growth and development. Further analyses are required to better understand the significance of these data.

## Limits of the study

This study has several limitations that should be considered when interpreting the findings. First, the sample size was relatively small, which may limit the generalizability of the results and affect the stability of multivariate approaches such as sPLS. In addition, although peripheral blood transcriptomics provides valuable insight into the physiological state of the horses, it does not allow us to determine the specific tissues or cellular pathways driving the observed changes, nor does it establish causal relationships between gene expression and behaviour. Another important limitation concerns the short duration of the pasture exposure, which prevents us from assessing whether the beneficial effects observed in resilient horses persist over longer periods or whether the inflammatory signatures detected in non-resilient horses represent a transient adaptation phase. Finally, although sPLS is a powerful tool for identifying associations between behavioural traits and gene expression, the relationships uncovered remain correlational. Further studies with larger cohorts, longitudinal designs, and functional validation will be necessary to clarify the biological significance of these transcriptomic signatures and their role in behavioural resilience.

## Acknowledgements

We thank Claire Naylies and Yannick Lippi for their contribution to microarray fingerprints acquisition and microarray data analysis carried out at GeT Genopole Toulouse Midi-Pyrénées facility (https://doi.org/10.15454/1.5572370921303193E12).

## Data, scripts, code, and supplementary information availability

Data, script and supplementary tables and figures are available online: https://doi.org/10.5281/zenodo.18622341

Transcriptomic data are available online: https://www.ncbi.nlm.nih.gov/geo/query/acc.cgi?acc=GSE310717

