## Supplementary Tables and Figures for "Effects of a temporary period on pasture on the transcriptomic signature of horses housed in individual boxes"

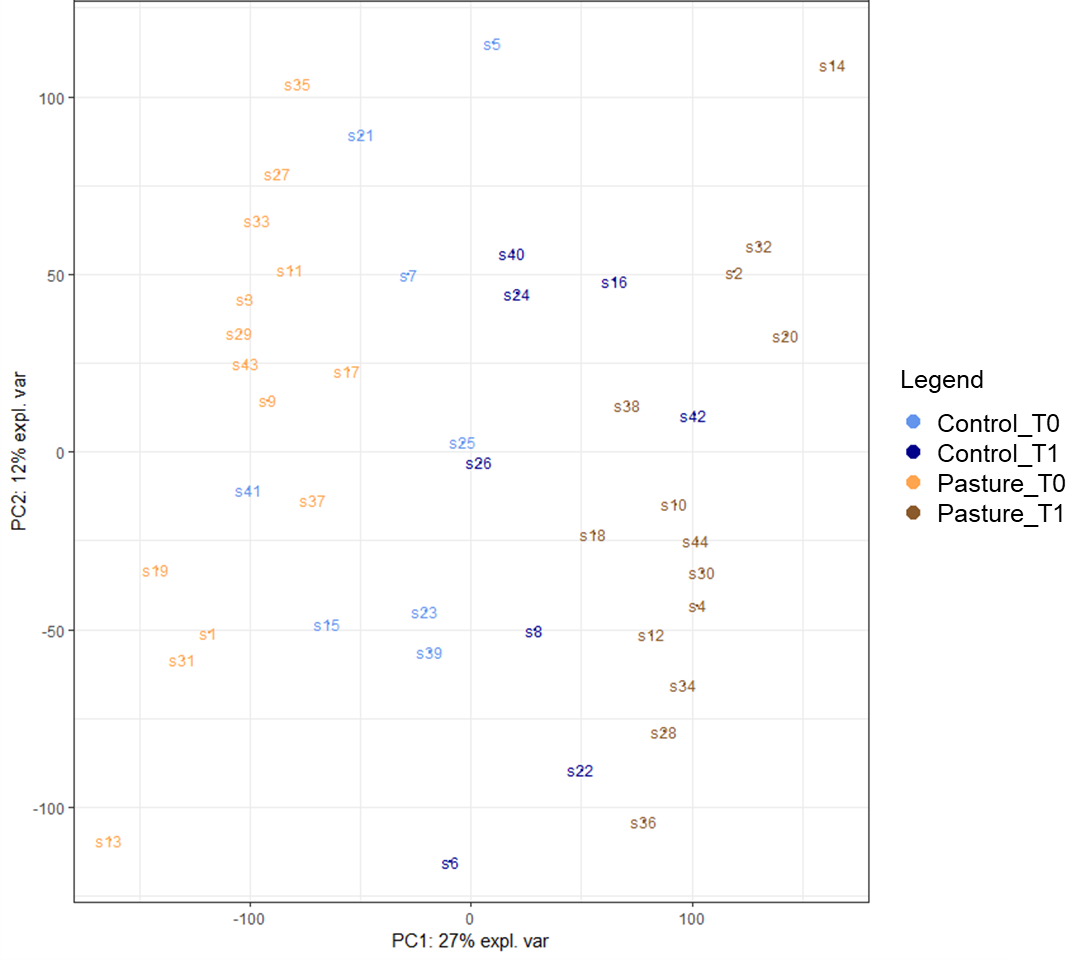

**Figure S1** – Sample plot from the PCA performed on the transcriptomic dataset. Samples are coloured by group: Control and Pasture horses before pasture (T0) and 3 months after returning to the box after the pasturing period (T1).

**Table S1** – Gene expression changes in cell proportions in the blood samples using the Celltype Computational Differential Estimation CellCODE R package: Control vs. Pasture horses before pasture (T0) and 3 months after returning to the box after the pasturing period (T1).

| **Blood cells** | **p value** | |
| --- | --- | --- |
|  | **T0** | **T1** |
| Neutrophil | 0,76 | 0,38 |
| Tcell | 0,38 | 0,45 |
| Monocyte | 0,71 | 0,11 |
| Bcell | 0,74 | 0,43 |
| NKcell | 0,45 | 0,22 |
| Megakaryocyte | 0,61 | 0,81 |
| Erythrocyte | 0,52 | 0,86 |

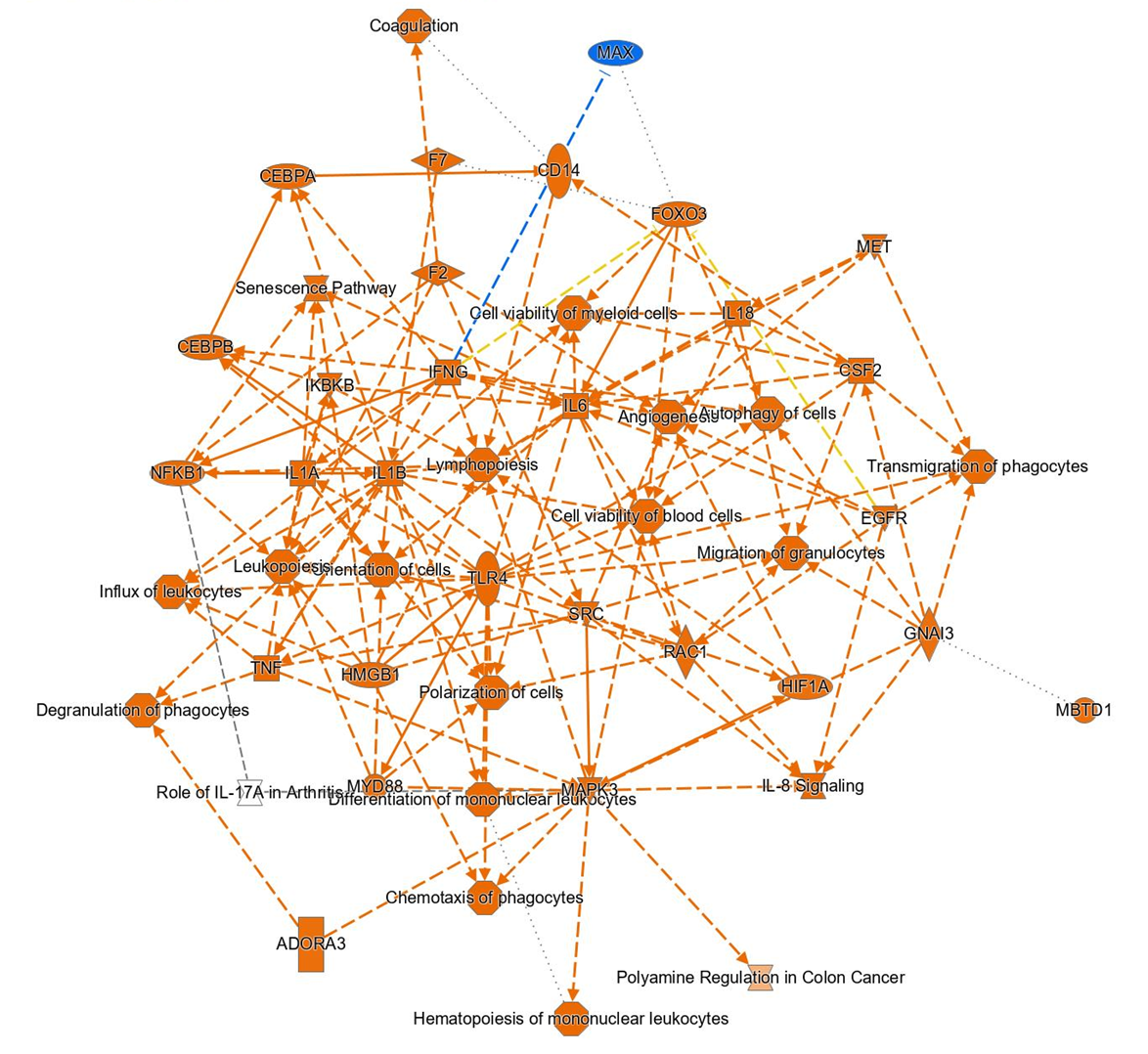

**Figure S2** – Graphical Summary of genes positively correlated to aggressiveness towards humans (r>0.5) generated through the use of QIAGEN IPA (QIAGEN Inc., <https://digitalinsights.qiagen.com/IPA>)

**Table S2** – Upstream regulators for the 414 genes positively correlated with aggressiveness towards humans in the sPLS analysis (r ≥ 0.5). Data were analysed with the use of QIAGEN IPA (QIAGEN Inc., <https://digitalinsights.qiagen.com/IPA>).

| **Regulator** | **Target genes** | **Pathway** | **p-value** |
| --- | --- | --- | --- |
| TGFB1 | AASS,ADCY9,AHR,ANXA11,ARF4,ARID5B,ATG4B,ATP13A3,BCL6,BTG1,CAB39,CCL3,CCL4,CD55,CDK2AP1,CHMP3,CHST11,CLIC4,CTNNB1,CXCL1,CXCL2,CXCL3,CXCL8,EGLN1,FCAR,FLI1,FNDC3B,FOSL2,FOXO3,FTH1,FTL,GADD45A,GCLC,GDPD5,GYG1,HBEGF,HCAR2,HEXIM1,HHEX,HMGB1,IDI1,IFRD1,KLF10,KLF3,LAMP2,LDLR,LIMK2,LITAF,MAP2K1,MAPK3,MAPKAPK2,MARCHF7,MDM2,NFKBIA,NUP153,OLR1,PDLIM7,PLAUR,PPP2R2A,PSME4,PTGS2,PXN,QKI,RAB18,RAB1A,RAB31,RAP1A,RARA,RBM3,SGK1,SMAD4,SOD2,SORD,STAT6,STK17A,TAB2,TLE4,TPM3,ZEB1 | Inflammation | 1,93E-11 |
| FSH | ACTR2,ADCY9,AHR,CD55,CTNNB1,FOSL2,GCLC,GNB1,KLF10,LDLR,MAP2K1,MAP4K4,PPP1CB,PPP1R2,PTGS2,PTP4A1,RAB1A,RAB27A,RAB31,RARA,SGK1,SMAD4,SNAP23,STK17A,STK24,TRIB1,UPP1,ZEB1 | Reproduction | 2,31E-09 |
| TP53 | AATF,AGO2,ASXL1,ATG4B,ATL3,BTG1,CCL4,CCNG1,CDKN2D,CLIC4,CLTC,CTNNB1,CXCL1,CXCL2,CXCL3,CXCL8,DMAC2L,EIF4G2,FOXO3,FTH1,G0S2,GADD45A,GDPD5,GRK3,GSK3B,HBEGF,HMGB1,LAMP2,LDLR,LIMK2,LYPLA1,MAFB,MAP2K1,MAPK3,MDM2,MED13L,MOCOS,MTDH,MYO5A,NFKBIA,NRF1,NUP153,PDHX,PDK3,PLAUR,PLPBP,PPP4R2,PTGS2,PTP4A1,RBM3,SAT1,SEC62,SESN2,SFN,SFPQ,SGK1,SNX2,SOD2,SON,STK17A,STOM,STRN3,TALDO1,TANK,TMEM43,TNFAIP8,TPM3,UBC,UPP1,VAMP3,VDAC1,WWP1,ZEB1 | Inflammation | 4,22E-09 |
| miR-124-3p | ABHD5,AHR,ATL3,CAPRIN2,CHSY1,CTDSP2,GNAI3,GSK3B,IQGAP1,ITPRID2,LDLR,LITAF,LMNB1,PDLIM7,PLSCR3,RASSF5,RFFL,STOM,TUBB6,VAMP3 | Tumor suppresor | 1,48E-08 |
| IL5 | BCL6,CCL3,CD55,CXCL8,DENND11,EGLN1,FOSL2,GADD45A,GCLM,HBEGF,HIGD1A,IDI1,LIMK2,LMNB1,MTDH,RBM3,RIPOR2,SFPQ,SGK1,SLAIN1,SNAP23,UPP1 | Inflammation | 2,30E-08 |
| CDK19 | ACVR1B,CCNG1,CXCL1,CXCL2,CXCL8,GADD45A,HBEGF,MDM2,MXD1,PLK3,SAT1,SFN,STOM,UPP1 | Growth/differenciation | 5,50E-08 |
| miR-155-5p | ANKFY1,CCL3,CCL4,CTNNB1,CXCL1,CXCL2,CXCL3,CXCL8,DCAF7,FOXO3,MAFB,MOSPD2,PICALM,PTGS2,TAB2,VAMP3 | Inflammation | 7,46E-08 |
| HNF4A | AASS,ACTR3,ADSS2,AP2A1,ASAH1,BCL6,BMP2K,BST1,BTG1,CCNG1,CD55,CHMP1B,CTNNB1,CUL2,CWC25,CXCL3,CXCL8,CYP26A1,DNAJC14,DYNC1LI1,FIG4,FOSL2,FTH1,G0S2,GHITM,GNAI3,GSK3B,GYG1,HBEGF,HMGB1,ITPRID2,IVNS1ABP,LDLR,MDM2,MKRN1,MOCOS,MTF1,N4BP1,NBR1,NDFIP1,NRAS,OSBP,OSBPL11,PCMT1,PITPNB,PJA2,PPP1R3B,PPP4R1,PPP6C,PRELID3B,PSMD10,PTGES3,RAB10,RAB18,RAB7A,RARA,RASSF5,SESN2,SGK1,SLC33A1,SMAD4,SNAP23,STIM1,STK24,STOM,TLE3,TM4SF4,TMEM43,TRAPPC8,UPF3B,UTP11,VDAC1,YKT6,ZNF281 | Cell structure | 1,19E-07 |
| **Regulator** | **Target genes** | **Pathway** | **p-value** |
| Lh | ACTR2,ADCY9,BCL6,CD55,CTNNB1,CXCL8,GNB1,LDLR,MAFB,MAP2K1,MAP4K4,PTGS2,PTP4A1,RAB1A,RAB27A,RAB31,SGK1,SNAP23,SOD2,STK17A,STK24,TRIB1,UPP1 | Reproduction | 1,53E-07 |
| CSF2 | CCL3,CCL4,CHST11,CLEC7A,CXCL1,CXCL2,CXCL3,CXCL8,GCLM,GRK3,HBEGF,HCAR2,HMGB1,IFRD1,LAMP2,MDM2,NFKBIA,OLR1,PLAUR,PTGS2,RARA,RBM3,RIPOR2,SGK1,SNAP23,SOD2,SORL1,UPP1,UTP11 | Inflammation | 2,42E-07 |
| ESR1 | AASS,AATF,ACTR2,ACVR1B,BCL6,CBL,CCL4,CCNG1,CCNI,CCPG1,CD55,CLEC7A,CTNNB1,CXCL1,CXCL3,CXCL8,CYFIP1,DR1,DUSP10,EIF4G2,FOSL2,GNB1,GYG1,HBEGF,IFRD1,IQGAP1,JAK1,LDLR,MARCKS,MDM2,NFKBIA,NRF1,NUP153,OLR1,PLAUR,PPP1R12A,PTGS2,RAB27A,RAB31,RARA,RGS3,RIT1,RP2,SFPQ,SGK1,SLC10A3,SNAP23,SON,SORD,SPAG9,STK3,TIRAP,TRIB1,YKT6 | Growth/differenciation | 2,70E-07 |
| FAS | CCL3,CCL4,CLTB,CXCL1,CXCL2,CXCL3,CXCL8,DUSP10,FCAR,FOXO3,LDLR,MAP4K4,MYO5A,NFKBIA,PDLIM7,PTP4A1,RARA,RGS3,SOD2,SORL1,STK17A,STK3,TRIB1 | Apoptose | 5,67E-07 |
| FOS | ATF7IP,CCNG1,CXCL3,CXCL8,DEDD,EIF4G2,FARS2,FOXO3,FTH1,GSK3B,LAMP2,MAP4K4,MDM2,MXD1,PLAUR,PPP2R2A,PTGS2,PTP4A1,PTS,PXK,QKI,RAB31,RARA,SNAP23,STAT6,STIM1,STK3,YWHAZ,ZEB1 | Apoptose | 1,39E-06 |
| NFKBIA | CDKN2D,CTNNB1,CXCL1,CXCL2,CXCL3,CXCL8,EIF4G2,GADD45A,LITAF,MDM2,NBR1,NFKBIA,NOCT,PICALM,PTGS2,RGS3,RNF19A,SAT1,SGK1,SMAD4,SOD2,SORL1,SYCP2,TNFAIP8,YWHAZ | Apoptose | 1,59E-06 |
| KRAS | ACTR3,AGO2,AHR,BPNT2,CAMKK2,CHSY1,CTNNB1,CXCL8,FTH1,GADD45A,GCLC,GCLM,HBEGF,HEXIM1,IVNS1ABP,JAK1,LAMP2,LMNB1,MAP2K1,MAPK3,MDM2,NFKBIA,NRAS,PAIP2,PARG,PTGS2,QKI,RAB27A,RAB7A,STIM1,STOM,TESC,TLE3,TRIB1,UPP1,ZEB1 | Inflammation | 1,69E-06 |
| CCL5 | AHR,CCL3,CCL4,CXCL2,CXCL3,CXCL8,MAPK3,OLR1,PLAUR,SGK1 | Inflammation | 1,79E-06 |
| IL15 | AHR,BTG1,CCL3,CCL4,CD55,CDK2AP1,CXCL2,CXCL8,EGLN1,GABPB1,HBEGF,HNRNPF,LYN,LYPLA1,MAP2K1,MAPK3,NFKBIA,PLEK,PPP6C,PTP4A1,SORL1,SRGN,SYAP1,TALDO1,UPP1 | Inflammation | 1,86E-06 |
| NDRG1 | CTNNB1,CXCL1,CXCL2,CXCL3,CXCL8,MDM2,SMAD4,ZEB1 | Apoptose | 2,36E-06 |
| NFE2L2 | AHR,CCL4,CXCL2,CXCL3,CXCL8,EIF4G2,ETV6,FOXO3,FTH1,FTL,GCLC,GCLM,IFRD1,LAMP2,NBR1,NOCT,PAFAH1B1,PTGS2,SAT1,SLC3A1,SOD2,SRGN,TALDO1,TMED2,UBC | Inflammation | 2,43E-06 |
| SELP | BST1,CCL3,CCL4,CXCL2,CXCL8,PLAUR,STAT6 | Cell structure | 2,85E-06 |
| RASSF5 | BZW1,CXCL2,CXCL8,PCMT1,RABIF,RAP1A,SAT1 | Tumor suppresor | 5,10E-06 |
| miR-130a-3p | CXCL2,CXCL3,MAFB,SMAD4,ZEB1 | Inflammation | 5,21E-06 |
| ERC1 | CXCL8,NFKBIA,PTGS2 | Inflammation | 6,11E-06 |
| CD3 | ACTR3,AHR,ATP11B,BCL6,BTG1,BZW1,CBL,CCL3,CCL4,CDK2AP1,CUL4A,CXCL8,FTL,GABPB1,HNRNPF,JAK1,LYN,LYPLA1,NFKBIA,PLEK,PTGS2,PTP4A1,RBM3,SORL1,SRGN,STIM1,STOM,STRN3,SYAP1 | Inflammation | 7,98E-06 |
| CISH | CCL4,CXCL3,HCAR2,IFRD1,OLR1,PLAUR,PTGS2,SORL1,STAT6 | Inflammation | 8,25E-06 |
| **Regulator** | **Target genes** | **Pathway** | **p-value** |
| RELA | AHR,ATP6AP2,CCL3,CXCL1,CXCL2,CXCL3,CXCL8,DOCK8,EGLN1,KLF10,LYN,MDM2,NFKBIA,OLR1,PLK3,PTGS2,PXN,QKI,RASSF5,SMAD4,SOD2,STIM1,TANK,ZBED4 | Inflammation | 9,49E-06 |
| CHD1 | CXCL1,CXCL2,CXCL3,PTGS2,SOD2 | Cell structure | 1,04E-05 |
| FN1 | ACTR2,CCL4,CCNI,CLIC4,CXCL1,CXCL2,CXCL3,CXCL8,MARCKS,NFKBIA,PLAUR,PXN,SNAP23,VDAC1,YWHAZ | Inflammation | 1,11E-05 |
| NFKB1 | AGO2,ATP6AP2,CCL3,CCL4,CXCL2,CXCL3,CXCL8,GADD45A,GNAI3,MDM2,NFKBIA,PLK3,PTGS2,PXN,RAB31,SOD2,STIM1,TANK | Inflammation | 1,11E-05 |
| SELPLG | BST1,CCL3,CCL4,CXCL2,CXCL8,PLAUR,STAT6 | Inflammation | 1,20E-05 |
| GNA12 | CXCL8,GCLC,GCLM,IQGAP1,JAK1,MAP2K1,PTGS2,PXN,RFFL | Inflammation | 1,42E-05 |
| ERK | CCL4,CTNNB1,CXCL1,CXCL2,CXCL8,FOSL2,FOXO3,GCLC,HBEGF,KLF10,LDLR,MAFB,MAP4K4,PTGS2,SFN,TNFAIP8 | Growth/differenciation | 1,44E-05 |
| EGR1 | CCL4,CDKN2D,CXCL2,CXCL3,CXCL8,FTL,GADD45A,HBEGF,LDLR,MXD1,PTGS2,SGK1,SOD2,STIM1 | Growth/differenciation | 1,48E-05 |
| Ige | BCL6,CCL3,CCL4,CHST11,CLEC7A,CXCL2,CXCL3,CXCL8,EIF4G2,HBEGF,MAPKAPK2,NFKBIA,PLK3,PPP1R3B,PTGS2,RAB7A,TPM3,TUBB6,YKT6 | Inflammation | 1,54E-05 |
| SETD2 | CTNNB1,CUL2,FOXO3,GSK3B,HHEX,QKI,RTN4 | Cell cycle | 1,64E-05 |
| LTB4R | CLEC7A,CXCL2,CXCL3,CXCL8,LDLR,PTGS2 | Inflammation | 2,21E-05 |
| C5 | BCL6,CCL3,CCL4,CD55,CXCL1,CXCL2,CXCL3,CXCL8,NFKBIA,PLK3,RGS3 | Inflammation | 2,28E-05 |
| FBXO32 | CXCL2,CXCL3,GADD45A,GCLC,ISYNA1,PTGS2,RAB27A,SLC33A1,SOD2 | Growth/differenciation | 2,34E-05 |
| ITGAX | CCL3,CCL4,CXCL8 | Inflammation | 2,41E-05 |
| TGIF1 | CCL3,CXCL1,CXCL2,CXCL3,CXCL8 | Growth/differenciation | 2,49E-05 |
| IgG | AHR,ASAH1,ATP1B3,CCL3,CCL4,CXCL8,DENND5A,FOSL2,HMGB1,LDLR,MAFB,MXD1,NFKBIA,PLAUR,PTGS2,SGK1,STAT6,STK24 | Inflammation | 2,99E-05 |
| NFkB (complex) | AHR,ATP6AP2,CCL3,CCL4,CTNNB1,CXCL1,CXCL2,CXCL3,CXCL8,FTH1,G0S2,GADD45A,GCLC,KLF3,LAMP2,LITAF,MDM2,MFHAS1,NFKBIA,OLR1,PLAUR,PLK3,PTGS2,PXN,SLC3A1,SOD2,STIM1,TNFAIP8 | Inflammation | 3,09E-05 |
| IKBKB | CCL3,CCL4,CTNNB1,CXCL1,CXCL2,CXCL3,CXCL8,EGLN1,FOXO3,GADD45A,LMNB1,MDM2,NFKBIA,NOCT,PTGS2,SGK1,SOD2 | Inflammation | 3,09E-05 |
| GNA13 | CXCL1,CXCL2,CXCL3,CXCL8,PTGS2 | Cell structure | 3,22E-05 |
| HMGB1 | CCL3,CCL4,CCNG1,CXCL2,CXCL3,CXCL8,HMGB1,MDM2,PTGS2 | Inflammation | 4,45E-05 |
| TNFSF11 | AHR,BST1,CCL4,CTNNB1,CUL4A,CXCL3,CXCL8,GCLC,IFRD1,MAFB,MAP2K1,NFKBIA,PLAUR,PTGS2,SOD2,YWHAZ | Apoptose | 4,45E-05 |
| **Regulator** | **Target genes** | **Pathway** | **p-value** |
| UPF1 | IQGAP1,LMNB1,NRAS,SGK1,SMG7,UPF3B | Regulator Of Nonsense Transcripts | 4,47E-05 |
| GNA14 | CXCL3,CXCL8,GADD45A,HBEGF,OLR1,PLAUR,PTGS2,SGK1 | Cell cycle | 5,11E-05 |
| mir-122 | CCNG1,CLIC4,FOSL2,GNPNAT1,IQGAP1,PXN,RBM3,SCAF11,VAMP3 | Mitochondrial metabolism | 5,29E-05 |
| LDL | CCL3,CCL4,CTNNB1,CXCL1,CXCL2,CXCL3,CXCL8,G0S2,GCLC,GCLM,LDLR,OLR1,PTGS2,RARA,SOD2,TANK | Cholesterol | 5,30E-05 |
| CD40LG | AHR,BCL6,BTG1,CCL3,CCL4,CCNG1,CXCL1,CXCL2,CXCL8,FOSL2,G0S2,GADD45A,MARCKS,NFKBIA,PLAUR,PLEK,PTGS2,SMG7,SOD2,TANK | Inflammation | 5,36E-05 |
| Jnk | ASAH1,BTG1,CCL3,CCL4,CD55,CXCL3,CXCL8,FLI1,GADD45A,PLAUR,PTGS2,RARA,SESN2,SOD2 | Inflammation | 5,38E-05 |
| HGF | AHR,BMP2K,CCL4,CCNG1,CTNNB1,CUL2,CXCL2,CXCL8,HBEGF,LDLR,LGMN,MAPKAPK2,MDM2,PLAUR,PPP2R2A,PTGS2,SGK1,SLK,SOD2,TRIB1,TUBGCP3,YWHAZ,ZEB1 | Growth/differenciation | 5,84E-05 |
| MARCHF3 | CXCL1,CXCL8,NFKBIA | Transport | 5,95E-05 |
| miR-145-5p | FBXO28,FLI1,LAMP2,MDM2,MYO5A,RAB27A,SOD2,TPM3 | Unknown | 6,24E-05 |
| JUN | BTG1,CXCL1,CXCL3,CXCL8,FOXO3,FTH1,GADD45A,GCLC,GSK3B,LAMP2,MAPK3,MDM2,NFKBIA,PLAUR,PPP2R2A,PTGS2,RAB31,RARA,SGK1,SOD2,STAT6,STIM1,ZEB1 | Inflammation | 6,39E-05 |
| BTG2 | CXCL1,CXCL2,CXCL3,CXCL8,RARA,SOD2 | Cell cycle | 7,15E-05 |
| NPM1 | CCL3,CCL4,CTNNB1,CXCL2,CXCL3,SOD2 | Inflammation | 7,15E-05 |
| MEOX2 | CDKN2D,CXCL1,CXCL2,CXCL3,CXCL8,HBEGF | Growth/differenciation | 7,15E-05 |
| PF4 | CCL3,CCL4,CXCL3,CXCL8,FLI1,NFKBIA,RARA,SOD2 | Inflammation | 7,56E-05 |
| CITED2 | ABTB2,CUL2,CXCL1,CXCL2,CXCL3,CXCL8,DR1,HCAR2,IFRD1,MARCHF5,MXD1,NRAS,SAT1,UPP1 | Growth/differenciation | 7,56E-05 |
| IL2 | ACVR1B,ADCY9,AHR,ARHGEF18,BCL6,CCL3,CCL4,CCNG1,CXCL3,CXCL8,FOXO3,HBEGF,HHEX,IDI1,IFRD1,JAK1,KLF3,LDLR,MAP2K1,MAPKAPK2,PPP6C,PTGS2,RFFL,SNAP23,SORD,STK17A,TNFAIP8,UPP1,VDAC1 | Inflammation | 7,74E-05 |
| NfkB-RelA | CCL3,CXCL3,CXCL8,NFKBIA,PTGS2 | Inflammation | 7,92E-05 |
| LRPAP1 | CXCL8,LDLR,MAPK3,PTGS2,SORL1 | cholesterol | 7,92E-05 |
| IL32 | CCL3,CCL4,CTNNB1,CXCL1,CXCL3,CXCL8,PTGS2 | Inflammation | 7,93E-05 |
| RB1 | CLIC4,CXCL2,CXCL3,CXCL8,GADD45A,GHITM,GPT2,HSBP1,KLF10,MAPK3,MCMBP,MFN2,MXD1,NFKBIA,NRF1,OLR1,PBX3,POLD3,RAB27A,SOD2,STK3,VDAC1,ZEB1 | Tumor suppresor | 8,84E-05 |
| F7 | CXCL2,CXCL8,GADD45A,HBEGF,PLAUR,PTGS2 | Coagulation | 9,55E-05 |
| PLCE1 | CXCL2,CXCL3,CXCL8,PTGS2 | Growth/differenciation | 9,62E-05 |
| **Regulator** | **Target genes** | **Pathway** | **p-value** |
| mir-375 | CXCL2,MTDH,QKI,YWHAZ | Unknown | 9,62E-05 |
| APP | ARF4,ATF1,BASP1,CCL3,CCL4,CLTB,CLTC,CTNNB1,CXCL1,CXCL2,CXCL3,CXCL8,CYFIP1,FOXO3,FTH1,GNB1,GSK3B,HCAR2,HMGB1,LDLR,MAFB,MAPKAPK2,NBR1,OLR1,PTGS2,RBM3,SEPTIN7,SOD2,STAT6,TPM3,TUBB6,UBC,VAMP3,VDAC1,YWHAZ | Growth/differenciation | 9,84E-05 |

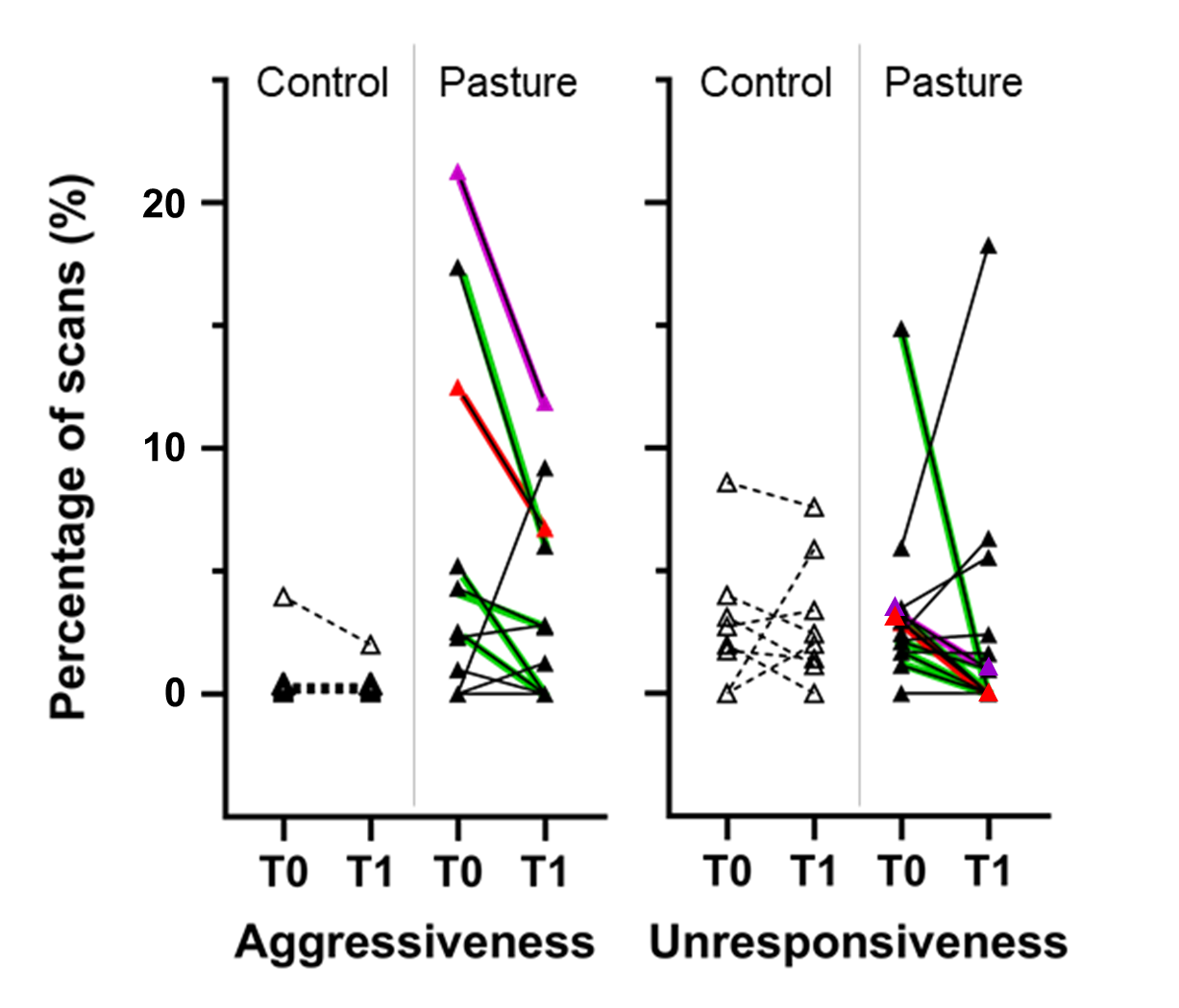

**Figure S3** – Percentage of scans recorded for aggressive behaviours towards humans and unresponsiveness to the environment before pasture (T0) and 3 months after returning to the box after the pasturing period (T1) in the Control (n=8) and Pasture groups (n=14). For the Pasture group, resilient horses are colored in green. The red and purple horses are resilient to both aggressiveness and unresponsiveness.

Control

Pasture
